# Lamins promote trophoblast lineage-associated transcription and trophoblast giant cell development in placental organogenesis

**DOI:** 10.64898/2026.08.21.746365

**Authors:** Sara Debic, Jiabiao Hu, Xiaobin Zheng, Yixian Zheng

## Abstract

Lamins are the major structural components of the nuclear lamina with a variety of roles in development and organogenesis. However, lamins’ function during trophoblast development, the first lineage to differentiate during mouse embryogenesis, remains unexplored. By utilizing an *in vitro* trophoblast stem cell differentiation model in a lamin null setting, we uncover that lamins maintain expression of genes related to trophoblast differentiation, while repressing genes involved in maintaining trophoblast stem cell stemness and off-lineage development. By deleting different combinations of lamins in mice, we show that both lamin triple-knockout and lamin-A and -B1 (lamin-A/B1) double-knockout result in placental defects, including reduced placenta size and disrupted placental organogenesis at embryonic day (E)9.5. At this stage, lamin-A/B1 are expressed in trophoblast giant cells of the placenta, and lamin-A/B1 loss leads to their impaired maturation *in vivo*. Lamin-A/B1 double knockout trophoblast giant cells exhibit reduced nuclear size along with a reduction of DNA damage signaling foci, suggesting a role for lamins in supporting trophoblast giant cell polyploidization. Similar to the transcriptional dysregulation observed during differentiation of lamin triple knockout trophoblast stem cells *in vitro*, lamin-A/B1 knockout *in vivo* results in downregulation of genes related to trophoblast giant cell function and upregulation of off-lineage genes. Our results suggest lamins are required for placental organogenesis by maintaining polyploidization and lineage-associated transcriptional programs in trophoblast giant cells.

**Highlights:**

- Lamins regulate lineage-associated transcription in a trophoblast stem cell model
- Lamin-A and -B1 support placental organogenesis
- Lamin-A and -B1 promote maturation of trophoblast giant cells (TGCs)
- Lamin-A and -B1 maintain expression of TGC genes relevant for placental function

## Introduction

Lamins are intermediate filament proteins that form the major structural component of the nuclear lamina, a proteinaceous layer found underneath the inner nuclear membrane [1]. Lamins interact with a wide variety of nuclear components, including inner nuclear envelope proteins, chromatin, and both the nuclear pore and linker-of-cytoskeleton and nucleo-skeleton (LINC) complexes [1]. Mammals such as mice contain three lamin genes: *Lmnb1* and *Lmnb2* which encode for the B-type lamins designated lamin-B1 (LB1) and lamin-B2 (LB2), as well as *Lmna* which encodes for the A-type lamin isoforms lamin-A and -C (collectively referred to here as LA) [1]. Lamins exhibit a variety of roles during organismal development, which include supporting nuclear envelope integrity, preventing DNA damage, organizing nuclear pore and the Linker of Nucleoskeleton and Cytoskeleton (LINC) complexes, and maintaining transcriptional programs important for a given cell type [2], [3], [4], [5], [6], [7], [8], [9], [10], [11], [12], [13]. Lamins are thought to influence transcriptional programs by organizing lamina-<u>a</u>ssociated chromatin <u>d</u>omains (LADs) [14], [15], [16], [17], which are heterochromatic regions that interact with the nuclear lamina [18]. Due to lamins’ diverse roles and the presence of three lamin genes, it has been challenging to tease apart the unique and shared functions of individual lamin proteins in different tissues during development.

Knockout of either LB1 or LB2 and combined knockout of both LB1 and LB2 leads to perinatal death in mice associated with organ defects, including DNA damage and apoptosis in migrating neurons of the brain [2], [3]. By contrast, knockout of A-type lamins leads to death ∼2- 8 weeks after birth depending on the knockout model, with defects observed in the adipose tissues, hearts, and skeletal muscles [19], [20], [21]. While these studies reveal that lamins are required for proper development, whether the different lamins play redundant or unique roles in a given lineage during development remains poorly understood.

B-type lamins are expressed in all cell types throughout mouse development, while lamin-A expression is temporally restricted depending on the cell type [22], [23]. During the blastocyst stage of early mouse development, lamin-A expression is most prominent in trophoblast cells [24], which eventually contribute to the development of the placenta [24], [25]. By embryonic day(E) 8.5, lamin-A expression persists in trophoblast-derived cell types and is also detected in the extraembryonic yolk sac, while remaining absent from cells of the embryo proper [22], [23]. The yolk sac supports embryonic growth during early and mid-gestation [26], before the placenta takes over the majority of nutrient-supplying functions following the establishment of placental circulation around E10 [26]. As both A- and B-type lamins are expressed during trophoblast development, they may have important roles in this lineage, a possibility that has remained unexplored. Understanding how lamins support development of the trophoblast lineage during mouse embryogenesis would yield important insights into lamins’ fundamental roles during organogenesis.

The trophoblast lineage differentiates into multiple specialized cell types with important functions during embryogenesis. At the blastocyst stage, the trophoblast lineage consists of the mural trophectoderm lining the blastocyst cavity and the polar trophectoderm overlaying the inner cell mass [27]. During implantation, the mural trophectoderm differentiates into the primary trophoblast giant cells (TGCs), which are characterized as parietal TGCs as they line the implantation site and are important for the exchange of nutrients and endocrine signals [27]. By contrast, the polar trophectoderm continues to proliferate and eventually gives rise to all the remaining cell types of the trophoblast lineage, including spongiotrophoblast, glycogen trophoblast cells, several labyrinthine cell types, and a second wave of TGCs known as secondary TGCs [27]. The secondary TGCs are thought to arise from *Tpbpa*-positive trophoblast precursors and they differentiate into the four distinct subtypes of TGCs, including sinusoidal TGCs, canal TGCs, spiral-artery associated TGCs, and parietal TGCs, each with specialized roles at the fetal-maternal interface [27]. During terminal differentiation, the different TGC sub- types all undergo polyploidization of their genome through repeated rounds of DNA replication without intervening mitosis, a process known as endoreplication [27]. This process is thought to help TGCs support the massive transcriptional and biosynthetic output of proteins important for placental functions, including remodeling of the fetal-maternal interface, immunomodulation, and hormone synthesis [28]. Several studies have established roles for canonical cell cycle regulators, such as cyclins E1 and E2, as well as DNA damage signaling during TGC endoreplication [29], [30], [31]. However, much less is known about how genome amplification processes are coordinated within the cell nucleus during polyploidization. Given the established role of lamins in nuclear organization, we investigated the role of lamins in TGCs to gain insights. Furthermore, although endoreplication is a defining feature of TGC differentiation, it remains unclear how polyploidization is linked to the transcriptional programs that underlie mature TGC function. Studying lamins’ function in the context of TGCs and placental development may yield insights.

By studying the role of lamins in trophoblast stem cell differentiation *in vitro* and in placenta development *in vivo*, we discover lamins support transcriptional programs relevant for placenta development and function, including the maintenance of TGC transcriptional programs. Furthermore, our RNA sequencing and immunofluorescence studies suggest a role for lamin-A and -B1 in TGC differentiation and endoreplication, which may in turn support placental organogenesis and function.

## Results

### Lamin-triple knockout in trophoblast stem cells causes dysregulation of trophoblast gene expression programs

To investigate the role of lamins in trophoblast development, we used our previously generated lamin triple-knockout (lamin TKO) mouse embryonic stem cells (mESCs) [32]. We differentiated the lamin TKO and control mESCs containing two alleles of each lamin gene into trophoblast stem cells by overexpression of *Cdx2* according to published methods [33], [34].

Lamin TKO and control mESCs were electroporated with a plasmid containing a tamoxifen-inducible Cdx2-estrogen receptor (ER) fusion protein construct. Differentiation into trophoblast stem cells was then induced by culturing under feeder-free conditions in fibroblast-conditioned trophoblast stem cell media containing FGF4, heparin, and tamoxifen to induce Cdx2-ER (Figure 1A). After 6 days in these culture conditions, both lamin TKO and control cells formed flat, epithelial colonies characteristic of trophoblast stem cells.

**Figure 1.**
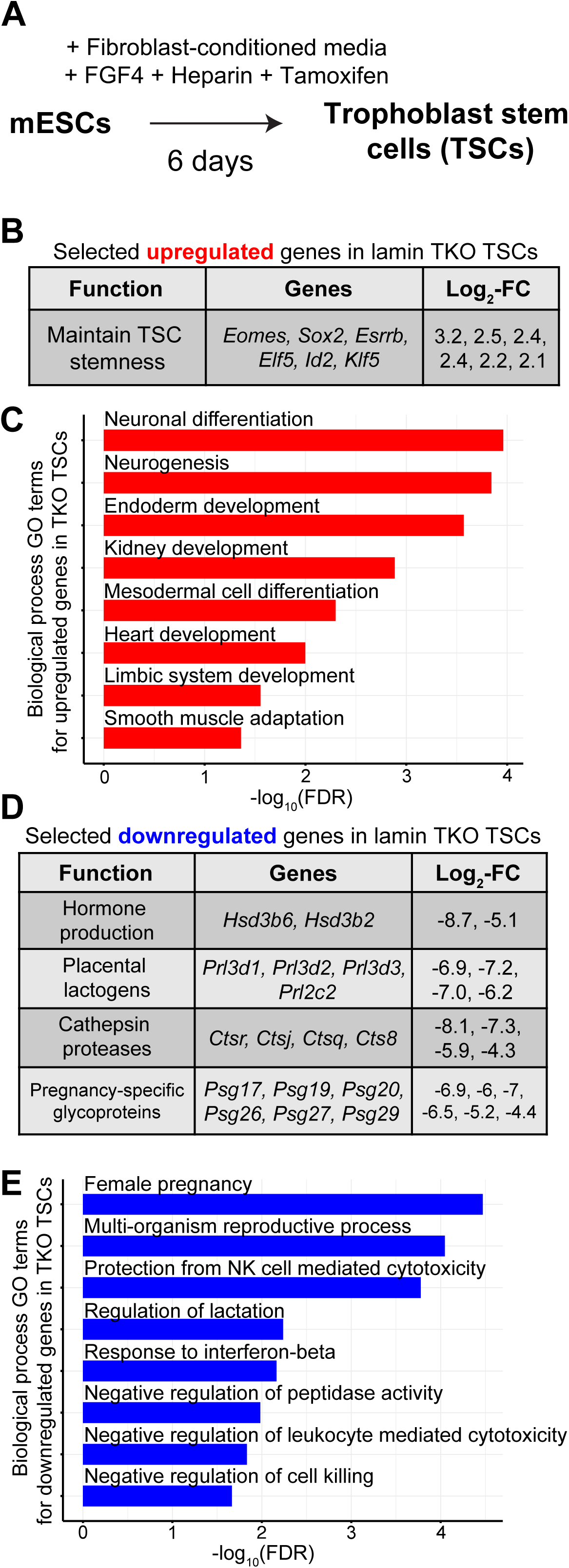
Lamin-triple knockout in a trophoblast stem cell differentiation model causes dysregulation of trophoblast and off-lineage gene expression. A) Schematic of the differentiation protocol used to generate trophoblast stem cells. B) Selected significantly upregulated genes in lamin TKO trophoblast stem cells with relevant trophoblast functions. C) Bar plot of selected GO terms associated with upregulated genes upon lamin TKO in trophoblast stem cells. D) Selected significantly downregulated genes in lamin TKO trophoblast stem cells with relevant trophoblast functions. E) Bar plot of selected GO terms associated with downregulated genes upon lamin TKO in trophoblast stem cells.

We next performed bulk RNA-sequencing on lamin TKO and control trophoblast stem cells to investigate transcriptional changes upon lamin loss. Using DESeq2, we found 551 significantly downregulated and 670 significantly upregulated genes (FDR < 0.05, log_2_-fold change cut-off of -2 or 2; Table S1). Among the upregulated genes were several transcription factors associated with maintaining trophoblast stem cell stemness and self-renewal, including *Eomes*, *Klf5*, *Elf5*, *Esrrb*, *Sox2*, and *Id2* [35], [36], [37], [38], [39], [40] (Figure 1B; Table S1). Gene Ontology (GO) term analysis of the upregulated genes revealed enrichment of GO terms related to the development of alternate organs and cell lineages, including neurons, endoderm, heart, and kidney (Figure 1C). Indeed, we found upregulation of the well-known endodermal transcription factor *Hnf1β* [41], as well as the characteristic skeletal muscle gene *Myoz1* [42] (Table S1). These findings indicate lamins promote the repression of off-lineage genes and genes related to maintaining stemness in trophoblast stem cells.

By contrast, genes downregulated upon lamin TKO in trophoblast stem cells are related to functions of differentiated trophoblast cells, and include genes characteristically expressed in trophoblast giant cells (TGCs). Notable examples of downregulated genes characteristic of TGCs include enzymes involved in placental progesterone synthesis, including *Hsd3b6* [43] (Figure 1D), and multiple placental lactogen genes such as *Prl3d1-3* (PL-1) and *Prl2c2* (Proliferin), which are hormones involved in maternal adaptation to pregnancy and placental angiogenesis [44], [45] (Figure 1D). We also observed reduced expression of cathepsin protease genes including *Ctsr*, *Ctsj*, *Ctsq*, and *Cts8* (Figure 1D), which are critical for trophoblast invasion and remodeling of the fetal-maternal interface [44], [46]. Lastly, numerous pregnancy-specific glycoprotein (PSG) genes were downregulated (Figure 1D); these contribute to immunomodulation and vascular remodeling essential for pregnancy maintenance [47], [48]. Consistent with these findings, GO term analysis of the downregulated genes in trophoblast stem cells revealed significant enrichment for processes related to pregnancy, lactation, and immune system regulation (Figure 1E). Together, these data show lamins promote the expression of genes relevant for trophoblast and placental function, including several genes characteristic of TGCs.

### Lamins are required for proper placental organogenesis

Given lamin TKO in an *in vitro* trophoblast stem cell differentiation model disrupts the expression of genes important for placental function, we next investigated whether lamin deletion leads to placental defects *in vivo* by using a whole-body lamin TKO mouse embryogenesis model. Since the trophectderm is the first lineage to differentiate, full body lamin TKO is likely to have major impact on trophoblast development during early embryogenesis and subsequent placenta development. We generated lamin TKO placentas by crossing males heterozygous for all three lamin genes and homozygous for Cre recombinase alleles driven by the beta-Actin promoter (*Actb-Cre^+/+^; Lmna^+/^*^Δ^*Lmnb1^+/-^Lmnb2^+/-^*) with females homozygous for floxed alleles of all three lamins (*Lmna^f/f^Lmnb1^f/f^Lmnb2^f/f^*). We focused on analyzing embryonic day (E)9.5 placentas since at this stage placentas contain clearly discernible spongiotrophoblast, labyrinth, and TGC layers, allowing us to assess potential defects in organogenesis caused by lamin loss. Hematoxylin and eosin staining of E9.5 sections showed that lamin TKO placentas were smaller and underdeveloped compared to lamin triple-heterozygote (*Lmna^+/^*^Δ^*Lmnb1^+/-^ Lmnb2^+/-^*) littermate controls (Figure 2A). Interestingly, similar placental defects were also observed in lamin-A/B1 double-knockout placentas retaining one copy of the lamin-B2 allele (*Lmna*^Δ*/*Δ^*Lmnb^-/-^Lmnb2^+/-^*) (Figure 2A). This data suggests loss of both lamin-A and -B1 is sufficient to disrupt placental organogenesis.

**Figure 2.**
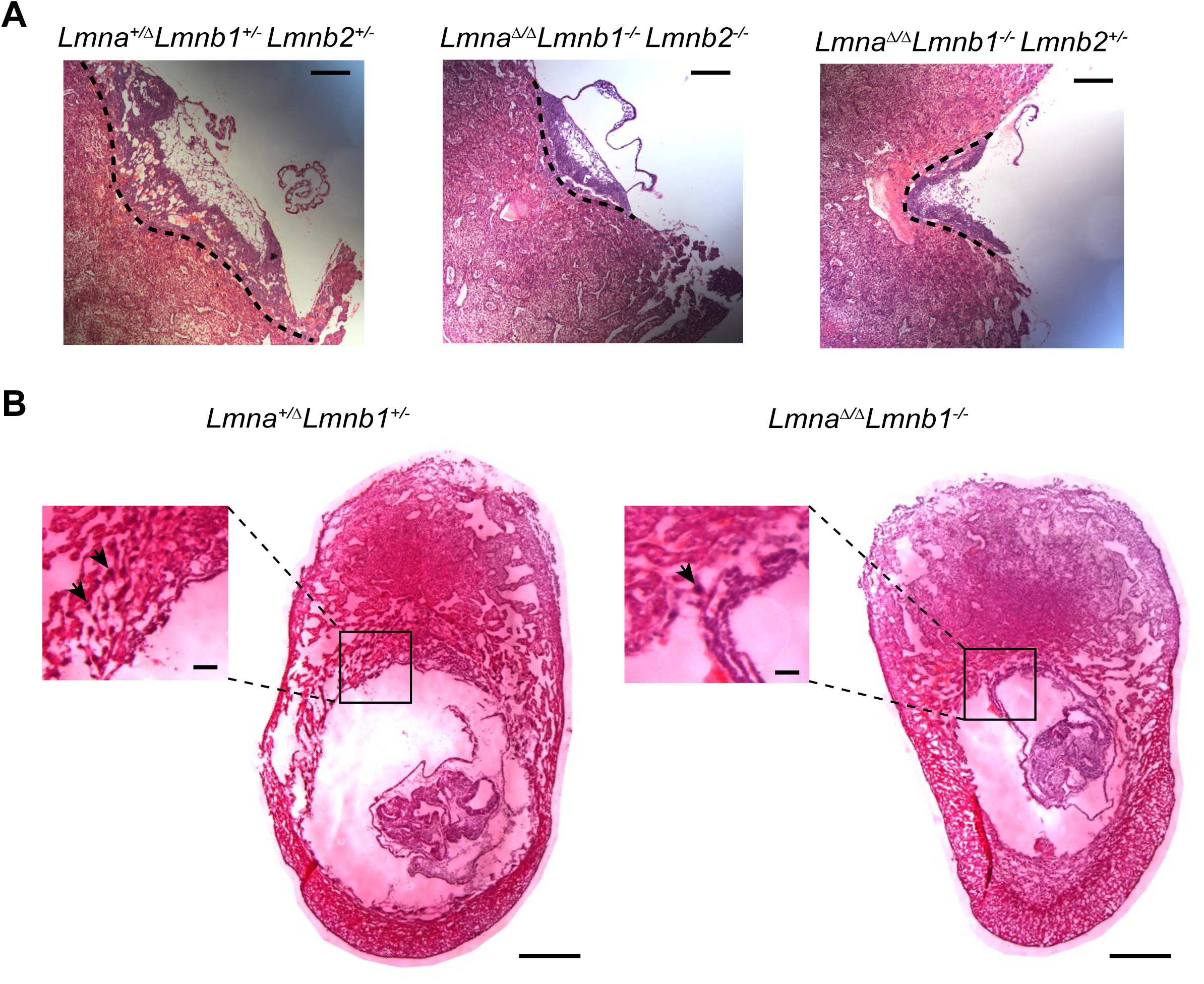
Disrupted organogenesis of lamin triple-knockout and lamin-A/B1 double-knockout placentas. A) Hematoxylin and eosin staining of *Lmna^+/-^ Lmnb1^+/-^ Lmnb2^+/-^*, *Lmna^-/-^ Lmnb1^-/-^ Lmnb2^-/-^*, and *Lmna^-/-^ Lmnb1^-/-^ Lmnb2^+/-^* placentas at embryonic day (E)9.5. Scale bar, 100 μm. B) Hematoxylin and eosin staining of *Lmna^+/-^ Lmnb1^+/-^* and *Lmna^-/-^ Lmnb1^-/-^* (lamin-A/B1 DKO) implantation sites and placentas. Scale bar, 500 μm. Zoom-ed in insets highlight the trophoblast giant cell layer, with individual giant cell nuclei indicated by black arrowheads. Scale bar of insets, 50 μm.

To determine whether the presence of only a single lamin-B2 allele contributes to the observed placental defects upon lamin-A/B1 loss, we next generated lamin-A/B1 double-knockout (DKO) placentas retaining two copies of the lamin-B2 allele by crossing *Actb-Cre^+/+^; Lmna^+/^*^Δ^*Lmnb1^+/-^*males with *Lmna*^f/f^*Lmnb1^f/f^* females. Hematoxylin and eosin staining of E9.5 implantation sites revealed that lamin-A/B1 DKO (*Lmna*^Δ*/*Δ^*Lmnb^-/-^Lmnb2^+/+^*) placentas exhibited reduced size compared to double-heterozygote (*Lmna^+/^*^Δ^*Lmnb1^+/-^Lmnb2^+/+^*) littermate controls (Figure 2B), closely resembling the phenotype of lamin TKO placentas. In addition, lamin-A/B1 DKO placentas showed a visibly thinner TGC layer (Figure 2B, see zoomed-in inset) and overall smaller implantation sites compared to controls (Figure 2B). Taken together, our results suggest both lamin-A and -B1, but not lamin-B2, play important roles to support proper placental organogenesis *in vivo*.

### Defects in trophoblast giant cells upon lamin-A and lamin-B1 loss in vivo

Given loss of both lamin-A and -B1 causes placental defects, we next examined which trophoblast cell types express both of these lamins at E9.5 so we could focus on these cells to investigate the underlying causes of disrupted placental organogenesis upon lamin-A/B1 DKO. Immunofluorescence staining of lamin-A and -B1 in control E9.5 tissue sections revealed both proteins are expressed in TGCs lining the implantation site and placenta (left panels in Figure 3A, 3B). We verified lamin-A and -B1 were not detected in TGCs of lamin-A/B1 DKO tissue sections (right panels in Figure 3A, 3B). These expression patterns suggest TGCs are a trophoblast cell type likely to be directly affected by lamin-A/B1 loss. As TGCs are important for mediating fetal-maternal interface remodeling and trophoblast invasion [27], defects in TGCs could underlie the reduced placental size and disrupted placental organogenesis observed upon lamin-A/B1 loss.

**Figure 3.**
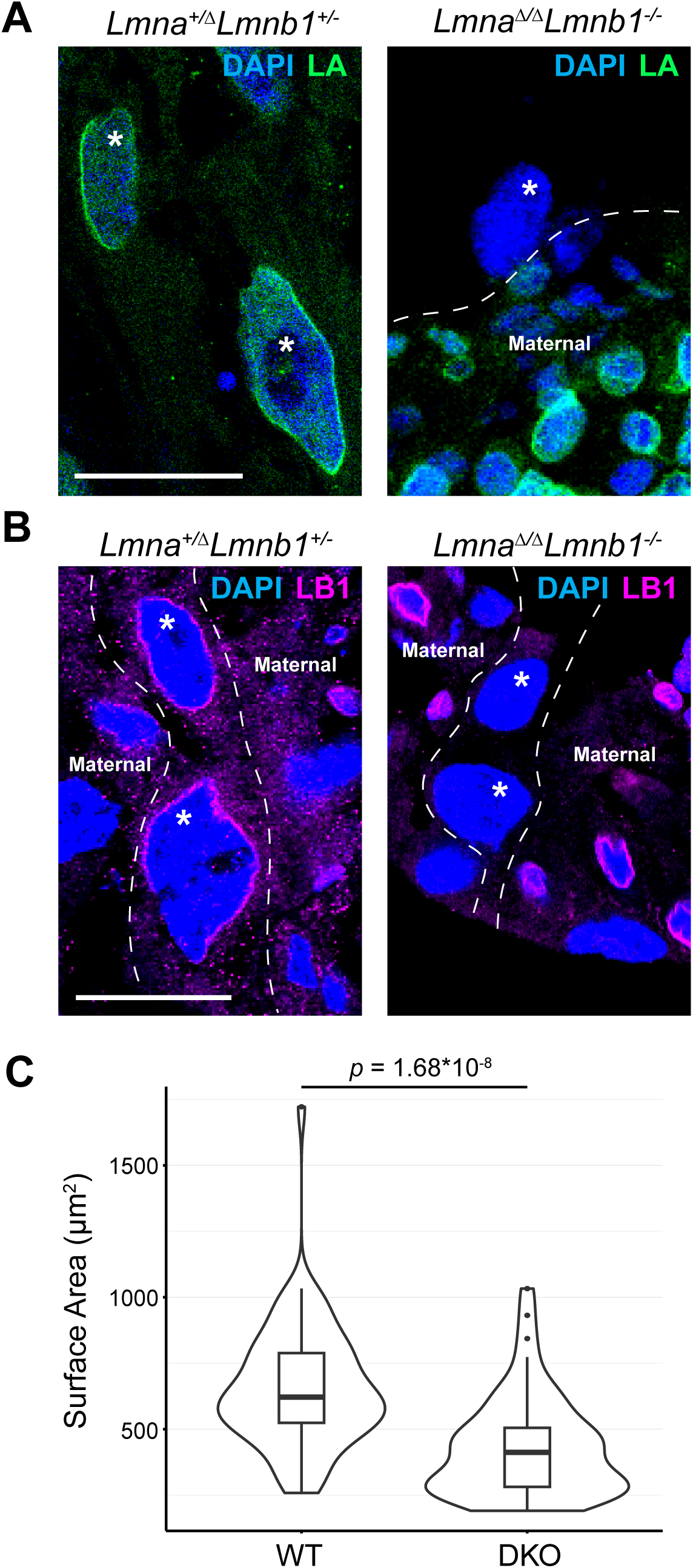
Lamin-A and -B1 maintain large size of trophoblast giant cell (TGC) nuclei. A) Representative immunofluorescence staining of lamin-A in *Lmna^+/-^ Lmnb1^+/-^* and *Lmna^-/-^ Lmnb1^-/-^* trophoblast giant cells (TGCs), which are labeled by white asterisks. Lamin-A positive cells in *Lmna^-/-^ Lmnb1^-/-^* sections are decidual cells of maternal origin. Scale bar, 20 μm. B) Representative immunofluorescence staining of lamin-B1 in *Lmna^+/-^ Lmnb1^+/-^* and *Lmna^-/-^ Lmnb1^-/-^* TGCs. Lamin-B1 positive cells in *Lmna^-/-^ Lmnb1^-/-^* sections are decidual cells of maternal origin. Scale bar, 20 μm. C) Violin plot showing the surface area of DAPI-stained TGCs from tissue sections. n = 2 biological replicates, 117 nuclei quantified; *p* = 1.68 * 10^-8^ (Wilcoxon rank-sum test).

We therefore next examined whether lamin-A/B1 loss causes defects in TGCs lining the implantation site and placenta. Interestingly, we observed nuclei of lamin-A/B1 DKO TGCs appeared smaller compared to control TGC nuclei (Figure 3A, 3B). To quantitatively assess differences in nuclear size, we measured the surface area of DAPI-stained control and lamin- A/B1 DKO TGC nuclei from 20 micron-thick tissue sections. We found lamin-A/B1 DKO TGC nuclei exhibited significantly decreased surface area compared to control nuclei (*p* = 1.68*10^-8^, Wilcoxon rank-sum test) (Figure 3C). Since nuclear enlargement accompanies terminal differentiation and polyploidization of TGCs [27], the reduced nuclear size of lamin-A/B1 DKO TGCs is consistent with altered TGC endoreplication and maturation.

To further investigate whether lamin-A/B1 loss affects processes associated with TGC maturation, we probed 53BP1 as a readout of DNA damage signaling associated with TGC endoreplication. Previous work has shown that TGC differentiation is accompanied by the accumulation of DNA damage signaling foci, presumably reflecting activation of DNA damage signaling from the accumulation of stalled replication forks and DNA damage during extensive DNA replication without intervening mitosis [31], [49]. Immunofluorescence staining of 53BP1 in TGCs from E9.5 tissue sections revealed decreased numbers of 53BP1 foci in lamin-A/B1 DKO TGCs compared to controls (Figure 4A, 4B) (*p* = 2.2*10^-16^, Wilcoxon rank-sum test). This result suggests that lamin-A/B1 loss is associated with reduced DNA damage signaling in TGCs, consistent with a role for lamin-A/B1 in supporting TGC endoreplication. Together, our *in vivo* findings demonstrate lamin-A/B1 maintain the large nuclear size and DNA damage signaling associated with polyploidization of TGCs, suggesting lamin-A/B1 support TGC endoreplication and differentiation.

**Figure 4.**
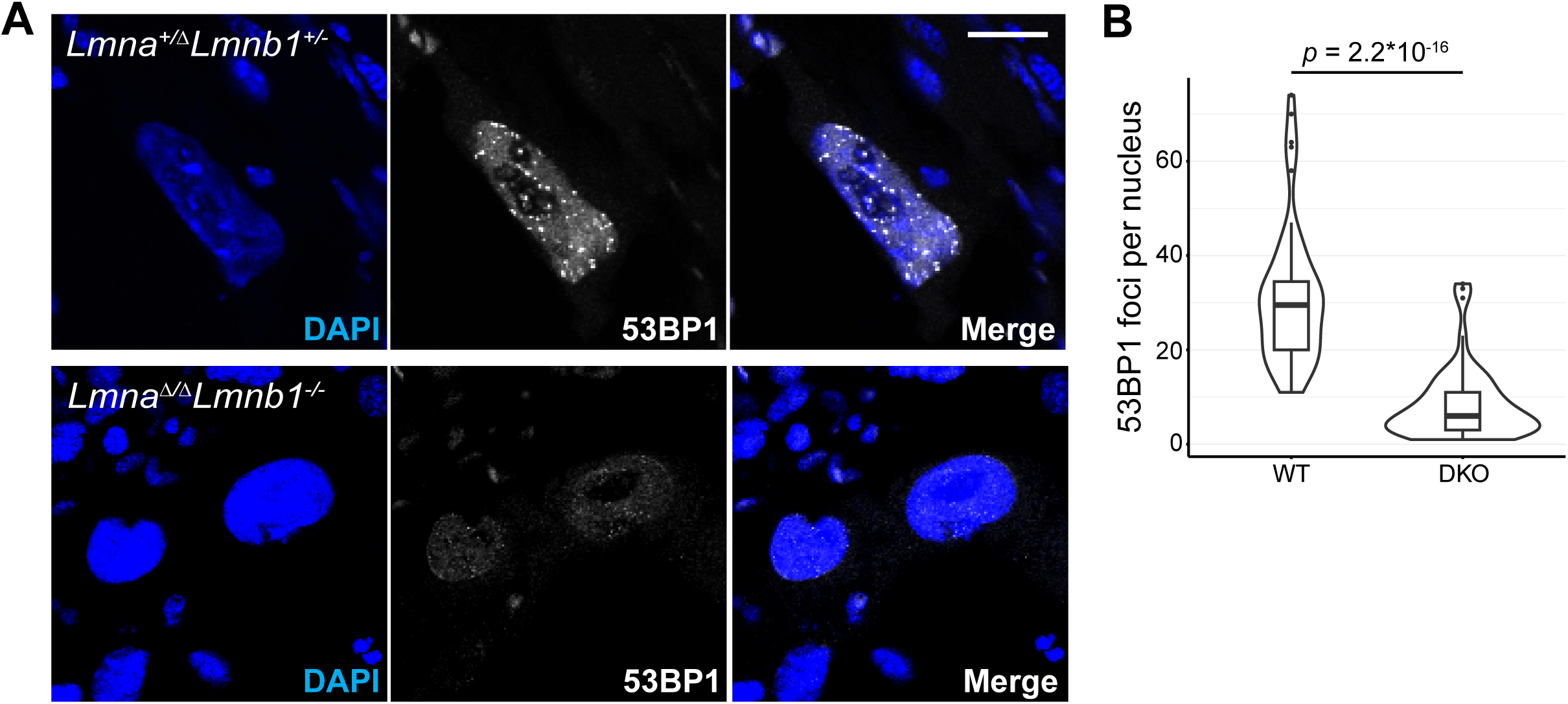
Decreased 53BP1 DNA damage signaling in TGCs upon lamin-A/B1 loss. A) Representative images of 53BP1 immunofluorescent staining in *Lmna^+/-^ Lmnb1^+/-^* and *Lmna^-/-^ Lmnb1^-/-^* TGCs. Scale bar, 20 μm. B) Violin plot showing the number of 53BP1 foci per TGC nucleus from tissue sections. n = 2 biological replicates, 117 nuclei quantified; *p* < 2.2 * 10^-16^ (Wilcoxon rank-sum test).

### Transcriptional changes of TGC-associated genes upon lamin-A/B1 loss in vivo

To further investigate how lamin-A/B1 loss impacts TGC differentiation, we assessed transcriptional changes upon lamin-A/B1 loss by RNA sequencing. Due to difficulty in isolating individual TGCs, we performed bulk RNA sequencing of whole lamin-A/B1 DKO and control placentas. Differential gene expression analysis of this dataset revealed 134 significantly downregulated and 67 significantly upregulated genes (FDR < 0.05, log_2_-fold change cut-off of - 2 or 2) (Table S2). Interestingly, several of these downregulated genes are normally expressed in TGCs. These include the prolactin gene *Prl3b1* and the cathepsin protease genes *Ctsr*, *Ctsj*, and *Ctsq* (Figure 5A, 5B, 5C), which were also downregulated in our lamin TKO trophoblast stem cell differentiation model (Figure 1D; Table S1). We also found downregulation of progesterone synthesis genes *Hsd3b2* and *Hsd3b3*, which are relevant to the known role of TGCs in progesterone production [50], as well as downregulation of the pregnancy-specific glycoprotein *Psg16* (Figure 5A; Table S2). Many additional cathepsin, pregnancy-specific glycoprotein, and progesterone synthesis genes were significantly downregulated (FDR < 0.05) in lamin-A/B1 DKO placentas but did not reach the log2-fold change cut-off of -2 (Table S2). The significantly downregulated genes in lamin-A/B1 DKO placentas are associated with GO terms related to placental and TGC functions such as hormone metabolism (Figure 5D), mirroring our findings using the *in vitro* trophoblast stem cell differentiation model. Furthermore, GO terms associated with significantly upregulated genes in lamin-A/B1 DKO placentas are related to off-lineage cell fates (Figure 5E; Table S2), also mirroring our findings in the *in vitro* trophoblast stem cell differentiation model. Overall, our results support the idea that lamins maintain polyploidization and transcriptional programs in TGCs to aid placental development.

**Figure 5.**
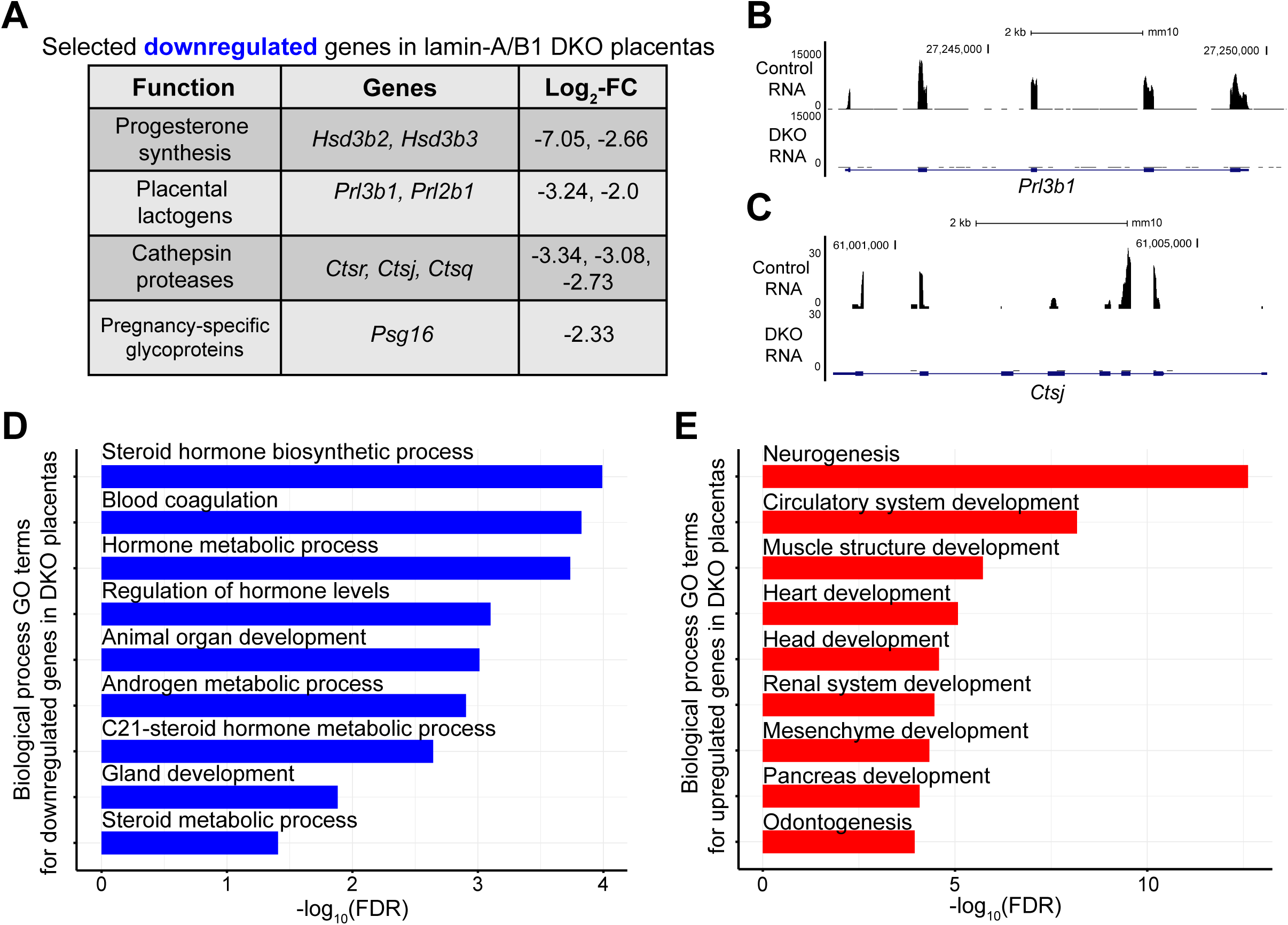
Transcriptional changes of lamin-A/B1 DKO placentas include trophoblast giant cell genes. A) Selected significantly downregulated genes in lamin-A/B1 DKO placentas with TGC-relevant functions. B) Genome browser view showing RNA tracks of *Prl3b1* and *Ctsj* expression from lamin-A/B1 DKO and control placentas. *Prl3b1* and *Ctsj* are genes known to be expressed in TGCs. D) Bar plot of selected GO terms associated with downregulated genes upon lamin-A/B1 DKO in placentas. E) Bar plot of selected GO terms associated with upregulated genes upon lamin-A/B1 DKO in placentas.

## Discussion

Previous developmental studies have emphasized the structural role of lamins in protecting migratory and mechanically stressed cell types from nuclear envelope rupture and DNA damage [4], [5], [6]. In this context, our *in vivo* findings from lamin-A/B1 DKO trophoblast are notable as we do not observe increased levels of 53BP1 DNA damage foci or overall grossly deformed nuclei in this lineage. Both our *in vitro* and *in vivo* results point to lineage-associated transcriptional defects as a major consequence of lamin depletion during trophoblast development. These findings are consistent with a growing body of evidence demonstrating lamins reinforce cell type-specific transcriptional programs across diverse cell lineages [7], [8], [9], [10], [11], [12]. Together, the evidence supports a model where lamins contribute to organogenesis not only by providing structural support to the nucleus, but also by maintaining specialized transcriptional programs required for lineage-relevant functions during development.

More specifically, our *in vivo* findings suggest lamin-A/B1 support polyploidization of TGCs and transcriptional programs associated with TGC functions. Previous studies have shown that certain gene clusters are selectively amplified during endoreplication of TGC genomes [28]. This is thought to help maintain the high expression levels of genes critical for placental function, analogous to the selective amplification of chorion genes in ovarian follicle cells of *Drosophila melanogaster* to support eggshell development [51]. Interestingly, we found that lamin loss both *in vivo* and *in vitro* leads to downregulation of key prolactin and cathepsin protease genes found within selectively amplified gene clusters [28]. These findings raise the possibility that lamin-A/B1 help sustain TGC transcriptional output by supporting the genome amplification programs on which selectively amplified, highly expressed TGC genes may depend on. Since some of these genes, including cathepsins *Ctsq* and *Ctsj*, are implicated in decidualization, trophoblast invasion, and vascularization [44], [46], transcriptional downregulation of highly expressed TGC genes provides a plausible explanation for the placental abnormalities observed *in vivo*. Thus, our data support a model where changes in TGC ploidy may contribute to changes in gene expression that ultimately affect placental development. Additional work is needed to more precisely define the changes in TGC ploidy upon lamin loss and to assess whether certain genomic regions or gene clusters are preferentially affected.

While our study suggests lamins support TGC polyploidization, the underlying mechanism remains unclear. We propose several, non-mutually exclusive possibilities. One possibility is that lamin loss affects endoreplication by disrupting expression of genes involved in DNA replication and DNA damage signaling, as has been described previously in multiple lamin knockout models [52], [53], [54]. A second possibility is that lamins may contribute more directly to the endoreplication process by recruiting factors that stabilize stalled DNA replication forks and coordinate DNA damage signaling that occurs during the extensive DNA replication that accompanies TGC polyploidization [31]. Indeed, several *in vitro* studies have demonstrated lamin-A protects stalled replication forks through recruitment of DNA repair proteins [55], [56], and both lamin A and -B1 have been implicated in the recruitment of 53BP1 to double-strand breaks [54], [57], [58], [59]. Thus, our observation that lamin-A/B1 DKO in TGCs leads to reduced 53BP1 foci is consistent with impaired 53BP1 recruitment and defective DNA damage signaling upon lamin loss. Impairment of DNA damage signaling and impaired resolution of stalled replication forks may disrupt the efficient endoreplication of TGC genomes. Future work utilizing models of trophoblast stem cell differentiation into TGCs should aid in investigating how lamins support the endoreplication processes of TGCs.

Interestingly, recent work in cell culture models has linked regulation of chromatin architecture by lamin-A/C to replication fork regulation under both basal conditions and replication stress [60], [61]. These findings raise the possibility that in TGCs, lamin-A/B1 may regulate chromatin architecture to support the extensive DNA replication processes required for polyploidization. For example, work in *Drosophila* has shown that chromatin decondensation at chorion gene loci is required for their developmental amplification, suggesting chromatin organization can influence DNA amplification and the resulting gene expression [62]. More broadly, recent *in vivo* work has suggested lamins maintain LADs and chromatin architecture to influence lineage-associated transcription factor function [9], [10], providing a unifying explanation for how lamin proteins support highly cell type-specific transcriptional programs during development. In TGCs, lamin-dependent chromatin organization may not only influence lineage-associated transcription factor function, but also contribute to the specialized genome amplification programs that support the high transcriptional demands of placental development. In the future, it will be interesting to study whether genomic regions that undergo selective amplification in TGCs are preferentially located near LADs, which could reveal a potential relationship between lamin-dependent chromatin organization and selective genome amplification.

In conclusion, our study supports a model where lamins contribute to organogenesis through the maintenance of lineage-associated transcriptional programs. By focusing on the trophoblast lineage, we find a specific role for lamin-A and -B1 in supporting TGC polyploidization and functional gene expression. Our work opens the door to further investigate how lamins maintain the unique 3D genome organization and nuclear processes of highly polyploid cells with specialized functions during organogenesis.

## Acknowledgements

We authors would like to thank Allison Pinder and Fred Tan for help with bulk RNA-sequencing; Mahmud Siddiqi for microscopy assistance; Lynne Hugendubler for technical assistance; Joseph Tran, Katherine Bossone, and Ross Pedersen for technical advice and helpful feedback; and additional members of the Carnegie Institution for Science and Zheng lab for helpful discussion. This work was supported by R01GM106023 (Y. Zheng, R.D. Goldman), R01GM110151 (Y. Zheng), and 1R01GM157598-01 (Y. Zheng).

## Author contributions

S.D. performed experiments in the lamin-A/B1 DKO mouse embryo model, bioinformatic analysis, visualization, and writing of the original draft. J.H. performed experiments in the *in vitro* trophoblast stem cell model and lamin TKO mouse embryo model. X.Z. advised on bioinformatic analyses. Y.Z. and S.D. performed writing and editing. Y.Z. supervised aspects of the work and acquired funding. Y.Z. and S.D. conceptualized the study.

## Declaration of AI

During preparation of the manuscript, S.D. utilized ChatGPT-5 mini (OpenAI) to assist with editing sections of the text for clarity and for finding references. S.D. also utilized Zo Computer (https://www.zo.computer) to aid literature survey and research the function of dysregulated genes. All content was reviewed, edited, and approved by the authors.

## Data availability

RNA-seq datasets generated in this study have been deposited in GEO under accession numbers GSE330009 and GSE330010. Code used to analyze RNA-seq data is publicly available at https://github.com/saradebic/trophoblast_transcriptional_analyses.

## Materials and Methods

### KEY RESOURCES TABLE

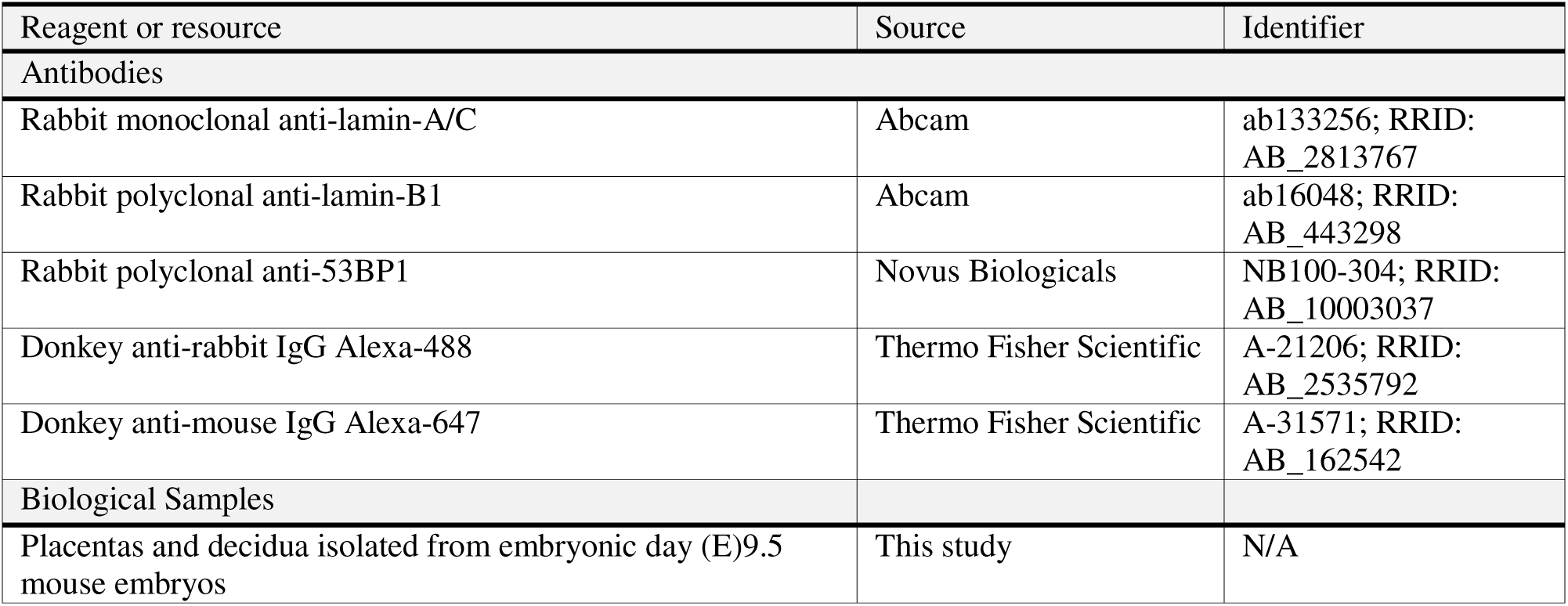
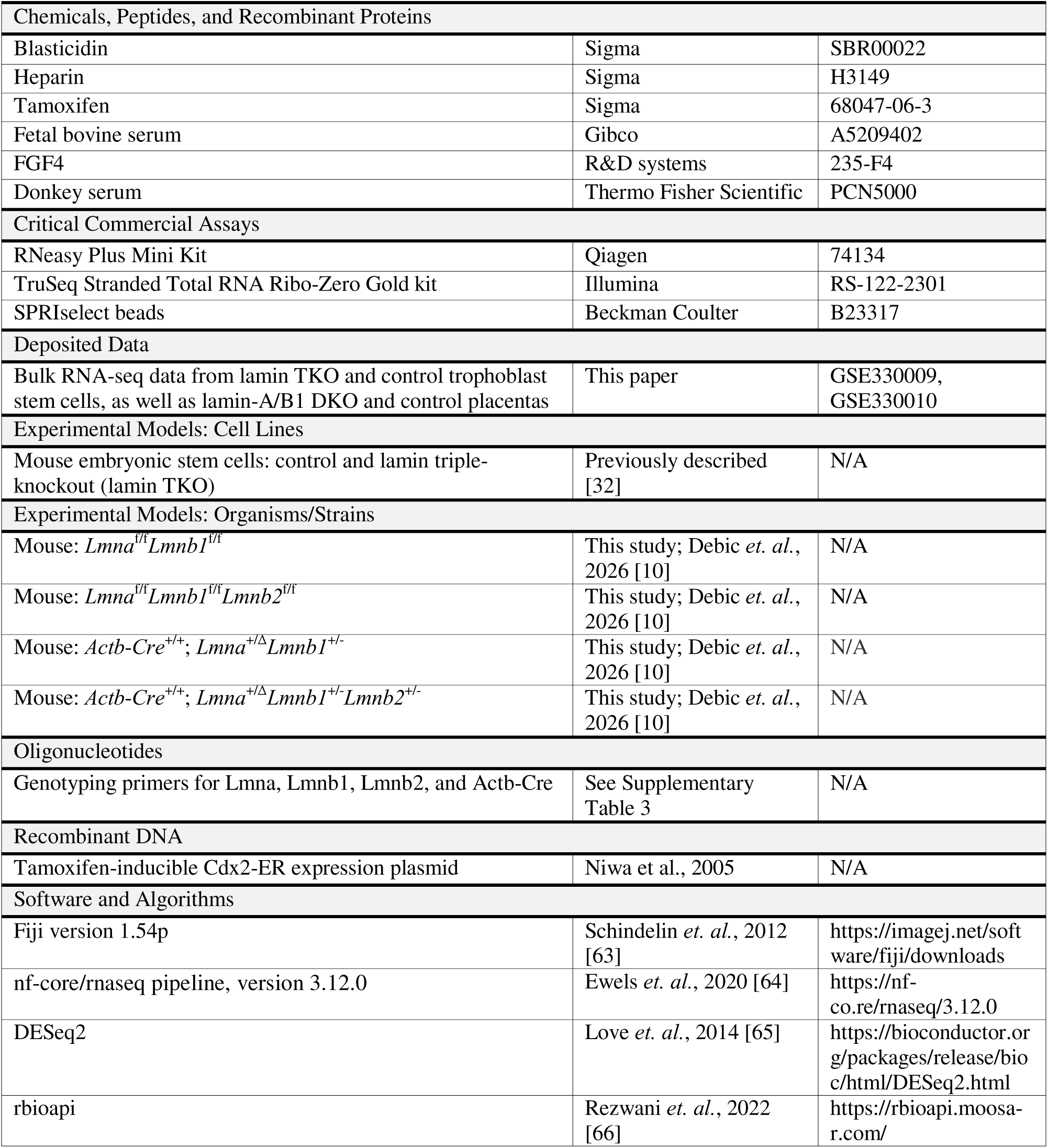

#### Trophoblast stem cell culture and differentiation

Mouse embryonic stem cells were electroporated with a plasmid containing the previously published Cdx2-ER construct [33] and a blasticidin resistant cassette. Following selection with blasticidin, individual clones were isolated and stable cell lines containing the Cdx2-ER construct were established. To induce differentiation towards trophoblast stem cell-like cells (TSCs), the established mESC lines containing Cdx2-ER were plated on gelatin-coated plates at 1.2 * 10^4^ cells/cm^2^ under feeder-free conditions with media containing 30% basal TSC medium, 70% fibroblast-conditioned medium, 1 mg/mL heparin, and 1 mg/mL 4-hydroxy-tamoxifen for Cdx2-ER activation. The cells were cultured under these conditions for 6 days, changing media every 2 days, to create trophoblast stem cell-like cells. For further differentiation, the trophoblast stem cell-like cells were cultured for an additional 6 days in 100% basal TSC medium. Basal trophoblast stem cell medium consisted of RPMI 1640 (Thermo Fisher, Cat# 21870-092) supplemented with 20% fetal bovine serum (FBS), 1x sodium pyruvate, 1x GlutaMAX, 0.1 mM β-mercaptoethanol, and 1x penicillin–streptomycin. Immediately prior to use, basal trophoblast stem cell medium was supplemented with 25 ng/mL FGF4 (PeproTech; prepared as a 1000x stock in PBS containing 0.1% BSA) and 1 µg/mL heparin. Fibroblast-conditioned medium was prepared using mouse embryonic fibroblast (MEF) feeder cells cultured in MEF medium consisting of KnockOut DMEM (Invitrogen, Cat# 10829-018) supplemented with 15% FBS (Invitrogen, Cat# 26140-079), 1× non-essential amino acids (NEAA; Invitrogen, Cat# 11140- 050), and 1× penicillin–streptomycin (Invitrogen, Cat# 15140-122). MEFs were seeded in 10 cm dishes (day 0), and the medium was replaced with 10 mL basal TSC medium on day 1. Conditioned medium was collected on day 4 and replaced with fresh basal trophoblast stem cell medium; this collection step was repeated twice. Collected media were pooled, filtered, aliquoted, and stored at −20 °C.

#### Mouse husbandry

All experimental mice were maintained in a controlled facility under a 12 h light/dark cycle at 20–22 °C and 30–50% humidity. Bedding, food, and water were changed routinely. Animal health was monitored via a sentinel surveillance program in which CD1 sentinel mice were exposed to soiled bedding from colony cages and screened quarterly for viral and parasitic pathogens by serology and PCR (Charles River Laboratories). All animal procedures were approved by the Institutional Animal Care and Use Committee (IACUC) and conducted in accordance with NIH and institutional guidelines. For timed matings, the date of vaginal plug detection was considered embryonic day (E)0.5.

#### Mouse strains

To generate lamin-A/B1 double-knockout (DKO) and lamin triple-knockout placentas, conditional alleles for *Lmna*, *Lmnb1*, and *Lmnb2* were used. A *Lmna*^f/f^*Lmnb1^f/f^* line was generated by crossing *Lmnb1^f/f^*mice carrying the *Lmnb1*^tm1a(EUCOMM)Wtsi^ allele (EUCOMM project, International Mouse Strain Resource) with *Lmna^f^*^/f^ mice (JAX stock #026284). For the lamin triple-flox line, the *Lmnb2^f/f^*allele was derived from *Lmnb2*^tm1a(KOMP)Wtsi^ (KOMP project, International Mouse Strain Resource) and the neomycin cassette flanked by FRT sites was removed by breeding with *Actb*-FLPe mice. *Actb-Cre^+/+^*; *Lmna^+^*^/Δ^*Lmnb1^+^*^/-^ and *Actb-Cre^+/+^*; *Lmna^+/^*^Δ^*Lmnb1^+^*^/-^*Lmnb2^+/-^*lines were generated by crossing mice heterozygous for *Lmna*, *Lmnb1,* and *Lmnb2* alleles generated previously [2], [21] (*Lmnb1* and *Lmnb2* deletion alleles available at Mutant Mouse Resource and Research Centers ID #42096) with mice homozygous for Cre recombinase driven by the human β-actin promoter (JAX stock #019099). Both lines were bred to homozygosity for the *Actb-Cre* allele. All mouse lines were maintained in a mixed genetic background of CD1, 129Sv, and C57BL/6J strains. Embryos of both sexes were used, and sex was not considered as a biological variable in this study.

#### Cryosectioning

Intact decidua were isolated at E9.5 and fixed in 4% paraformaldehyde (PFA) in PBS overnight at 4°C, followed by three washes in PBS. Next, the decidua were incubated in 30% sucrose solution in PBS overnight, and then embedded in OCT compound (Sakura, 4583) in cryomolds, and frozen at -80°C. Cryosectioning was performed using a Leica CM3050 S cryostat at −20°C. Decidua were sectioned into tissue slices ranging from 7 to 20 μm in thickness and mounted onto Superfrost Plus glass slides (VWR). These tissue slices included cross sections of the placenta and trophoblast giant cells. Genotyping was performed directly from tissue sections by scraping embryonic material into 0.1% Tween-20 in PBS, followed by lysis in RIPA buffer (50 mM Tris-HCl, 150 mM NaCl, 0.1% SDS, 0.5% sodium deoxycholate, 1% Triton-X-100, pH = 7.5) containing 40 μg of proteinase K. Samples were incubated overnight at 50°C with shaking. Genomic DNA was purified the following day using SPRIselect beads (Beckman Coulter) following the manufacturer’s protocol. 1 μL of supernatant was used as template DNA for PCR using the Terra PCR kit following the manufacturer’s protocol. PCR cycling conditions were as follows: 98°C for 2 minutes, followed by 35 cycles of 98°C for 10 seconds, 63°C for 15 seconds, and 68°C for 1 minute.

#### Hematoxylin and Eosin staining

Tissue sections were air-dried for several minutes to remove moisture. Hematoxylin staining was performed using 0.1% Meyer’s Hematoxylin (Sigma) for 10 min in a 50 mL conical tube. Slides were then rinsed in cool running distilled water for 5 min using a Coplin jar. Eosin staining was performed by briefly immersing slides in 0.5% eosin (1.5 g eosin dissolved in 300 mL 95% ethanol) 12 times. Slides were then rinsed in distilled water until eosin runoff was minimal, followed by sequential dehydration in 50% ethanol (10 dips), 70% ethanol (10 dips), 95% ethanol (30 seconds), and 100% ethanol (1 min). Slides were then cleared by briefly immersing in xylene several times. Finally, slides were mounted using ProLong Glass Antifade Mountant (Thermo Fisher, P36930), allowed to cure, and imaged using a Leica M125 dissecting microscope equipped with a Leica IC80 HD camera.

#### Immunofluorescence

Slides were permeabilized with 0.25% Triton-X-100 for 20 minutes at room temperature using a coplin jar and subsequently incubated overnight at 4°C in a humidified chamber with primary antibody diluted in block solution (10% donkey serum volume/volume, 10% BSA weight/volume, 10 mM sodium azide, and 0.1% Tween-20 in 1X PBS volume/volume). Immunofluorescence staining was performed using primary antibodies against lamin-A/C (Abcam, ab133256; diluted 1:200), lamin-B1 (Abcam, ab16048; diluted 1:200), and 53BP1 (Novus, NB100-304; 1:1000 dilution). The following day, samples were washed three times with PBS for 5 minutes each and incubated for one hour at room temperature protected from light with a 1:1000 dilution of Alexa Fluor-conjugated secondary antibodies (ThermoFisher) in block solution containing 5 μg/uL DAPI (ThermoFisher, D1306). After an additional PBS wash, the sections were mounted with ProLong Glass Antifade Mountant (ThermoFisher, P36930) and imaged using a Leica SP5 confocal microscope. Nuclear surface area was quantified using Fiji (ImageJ). DAPI-stained images were converted to 8-bit grayscale and thresholded using manual thresholding to generate binary masks of nuclei. Individual nuclei were identified using the “Analyze Particles” function with size and circularity filters set to exclude debris and smaller nuclei. For each segmented TGC nucleus, the “Area” measurement was extracted, and results were exported for downstream analysis using R statistical software. TGC nuclei were identified by their large size and characteristic positioning lining the implantation site and placenta, and only nuclei fully contained within the field of view were included in quantifications. To quantify 53BP1 foci, we used the Analyze Particles function in Fiji (ImageJ) after performing manual thresholding. 53BP1 foci were quantified per DAPI-stained TGC nucleus based on visual inspection of merged channels. Particles with a size between 0.1 and 100 μm^2^ and with a circularity between 0.3 and 1 were included, with identical thresholding and particle size parameters applied across all images.

#### Bulk RNA-sequencing

Total RNA was isolated from lamin TKO and control trophoblast stem cell cultures, as well as lamin-A/B1 DKO and control placentas, using the RNeasy Plus Mini kit (Qiagen, 74134) following the manufacturer’s protocol. Briefly, trophoblast stem cell cultures were washed with PBS and directly lysed in Buffer RLT Plus, while intact placentas were dissected from decidua, transferred to microcentrifuge tubes, and flash-frozen in liquid nitrogen for 5 seconds prior to lysis. Each placenta was then lysed in 350 μL of Buffer RLT Plus and homogenized by passing the lysate through a 20-gauge needle attached to a sterile syringe. Lysates from both trophoblast stem cell cultures and placental tissue were processed using the same column-based purification workflow according to the kit instructions, and RNA was eluted in 20 μL of RNase-free water. Ribosomal RNA was depleted using the TruSeq Stranded Total RNA Ribo-Zero Gold kit (Illumina, RS-122-2301). Libraries were sequenced on an Illumina NextSeq 500 platform using 50 bp single-end reads.

#### Bulk RNA-sequencing data analysis

Pre-processing of RNA-sequencing data was performed by running the nf-core/rnaseq pipeline version 3.12.0 using Nextflow version 22.10.1 [64]. Briefly, reads were mapped using the STAR aligner to the mouse genome mm10, and transcript quantification was performed using the Salmon software and GENCODE GTF file vM23. For differential gene expression analysis, we used the DESeq2 R package [65]. We used the merged gene counts file output from Salmon as input into DESeq2. We removed genes with a total RNA count less than 10 across all replicates from the differential gene expression analysis. We used FDR < 0.05 and a log_2_ fold-change cut- off of -2 or 2 as a threshold for differential gene expression. Biological process Gene Ontology (GO) term analysis of the differentially expressed genes was performed using the rbioapi R package [66].

## Supplementary Tables

Table S1. Significantly downregulated or upregulated genes in lamin TKO trophoblast stem cells, along with their associated GO terms.

Table S2. Significantly downregulated or upregulated genes in lamin-A/B1 placentas, along with their associated GO terms.

Table S3. PCR primers used for genotyping.

